# The antiviral potential of the antiandrogen enzalutamide and the viral-androgen interplay in seasonal coronaviruses

**DOI:** 10.1101/2023.11.25.568685

**Authors:** Oluwadamilola D Ogunjinmi, Tukur Abdullahi, Riaz-Ali Somji, Charlotte L Bevan, Wendy S Barclay, Nigel Temperton, Greg N Brooke, Efstathios S Giotis

**Affiliations:** School of Life Sciences, University of Essex, Colchester, UK; Department of Surgery and Cancer, Imperial College London, London, UK; Department of Infectious Diseases, Imperial College London, London, UK; Viral Pseudotype Unit, Medway School of Pharmacy, University of Kent, Chatham, UK

**Keywords:** Seasonal coronaviruses, antiandrogens, enzalutamide, TMPRSS2, androgen response element, SARS-CoV-2

## Abstract

The sex disparity in COVID-19 outcomes with males generally faring worse than females has been associated with the androgen-regulated expression of the protease TMPRSS2 and the cell receptor ACE2 in the lung and fueled interest in antiandrogens as potential antivirals. In this study, we explored enzalutamide, an antiandrogen used commonly against prostate cancer, as a potential antiviral against the human coronaviruses which cause seasonal respiratory infections (HCoV-NL63, -229E, and -OC43). Using lentivirus-pseudotyped and authentic HCoV, we report that enzalutamide reduced 229E and NL63 entry and replication in both TMPRSS2- and non-expressing immortalised cells, suggesting a TMPRSS2-independent mechanism. However, no effect was observed against OC43. To decipher this distinction, we performed RNA-sequencing analysis on 229E-and OC43- infected primary human airway cells. Our results show a significant induction of androgen-responsive genes by 229E compared to OC43 at 24 and 72h post-infection. The virus-mediated effect to AR signaling was further confirmed with a consensus androgen response element (ARE)-driven luciferase assay in androgen-depleted MRC-5 cells. Specifically, 229E induced luciferase reporter activity in the presence and absence of the synthetic androgen mibolerone, while OC43 inhibited induction. These findings highlight a complex interplay between viral infections and androgen signaling, offering insights for potential antiviral interventions.

## Introduction

Highly pathogenic coronaviruses, including SARS-CoV, MERS-CoV, and SARS-CoV-2, have caused deadly outbreaks in the 21st century.^1^ Four human CoV (HCoV: 229E, NL63, OC43, and HKU1) are endemic globally and cause 10-20% of seasonal upper respiratory (re-)infections in adults.^2^ HCoV- 229E and -NL63 are clustered phylogenetically within the genus Alphacoronavirus and emerged in humans from bat populations.^3^ Although these viruses are typically associated with common cold, they can cause severe pneumonia in immunocompromised individuals.^4,5^ NL63 is associated with croup^6^, 229E can cause respiratory distress in healthy adults sporadically^5^, HKU1 typically leads to mild respiratory symptoms, and OC43 is often implicated in severe respiratory infections, especially among vulnerable populations.^4,5^ There are currently no prophylactic vaccines or specific antiviral drugs approved for human use against HCoV. Interest in these viruses has been recently renewed as they can be handled in reduced biosafety laboratory containment thus providing an alternative to SARS-CoV-2 for preclinical screening and antiviral design. Furthermore, recent studies reported a possible cross-protective effect of preexisting HCoV-infection immunity on subsequent SARS-CoV-2 infection and on the severity of COVID-19 outcome.^7^

While there is no explicit report of sexual discordance in HCoV prevalence, it is becoming more evident that males of both younger and older ages are more susceptible to respiratory viruses in general and are also at a higher risk of severe disease outcomes when compared to females.^2^ The reasons are not entirely clear, but several explanations have been proposed, including immunological, genetic, hormonal and socio-behavioral factors.^8^ The sex discordance in COVID-19 outcomes with males generally faring worse than females has been associated with the primarily male hormones androgens and fueled interest in antiandrogens as potential antivirals.^8^ We previously demonstrated that enzalutamide, an antiandrogen typically used to prevent the androgen-mediated growth of castrate-resistant prostate cancers, reduces the expression of the transmembrane serine protease TMPRSS2, a key protease for SARS-CoV-2 cell entry, and thus holds potential as antiviral.^9^ TMPRSS2 is part of the type 2 transmembrane serine protease family and has been extensively studied in the context of prostate cancer metastasis. Its expression is regulated in response to androgens through direct transcriptional regulation by the androgen receptor (AR).^10^ Other studies showed that antiandrogens have pleiotropic effects *e.g.*, spironolactone can reduce the expression of the cell surface receptor of SARS-CoV-2 Angiotensin-Converting Enzyme 2 (ACE2) on the surface of lung cells decreasing therefore their susceptibility to the virus.^11^ Similar to TMPRSS2, ACE2 expression has also been shown to be regulated by the AR.^12^

The androgen regulation of TMPRSS2 and ACE2 raises the possibility that antiandrogens may inhibit the cell entry of other coronaviruses, which either enter target cells via a TMPRSS2-dependent pathway and/or use ACE2 as their cell receptor. Several cell receptors have been described for HCoV, including aminopeptidase N for 229E^13^, and ACE2 for NL63^14^. The cellular receptors for OC43 and the uncultivable in cell culture HKU1 are currently unknown but O-acetylated sialic acids, Kallikrein 13 and TMPRSS2 have been identified as important cell entry factors.^15,16^ Following binding to cell receptors, HCoV like most coronaviruses gain access to the cells either via direct fusion with the cell membrane or via cathepsin-mediated endocytosis.^17^ Several studies reported that circulating OC43, HKU1, and 229E HCoV generally use cell-surface TMPRSS2 for cell entry and not endosomal cathepsins in human airway epithelial cells.^18–20^ It was suggested that NL63 preferentially enters target cells through the endocytic route, but it can employ TMPRSS2 and bypass endocytosis in airway epithelial cells.^21^

In this study, we aimed to investigate the significance of TMPRSS2 in facilitating cell entry of NL63 and 229E viruses in A549 immortalised lung cells and evaluate the potential of the antiandrogen enzalutamide to inhibit virus entry. We observed reduced susceptibility to pseudotypes expressing the spike glycoproteins of NL63 and 229E viruses and wild-type NL63 and 229E viruses in enzalutamide-treated A549 human lung epithelial cells overexpressing ACE2. However, this effect was not observed for the OC43 wild-type virus. Collectively, our data suggest that antiandrogens are promising candidates for the development of broad-spectrum therapeutics to treat coronaviruses.

## Materials and methods

### Plasmids, cell lines, and inhibitors

The HCoV-229E and -NL63 seasonal S genes were synthesised in a pcDNA3.1^+^ backbone by GeneArt Gene Synthesis, Thermo Fisher. The human ACE2 receptor plasmid pCAGGS-ACE2 and the human TMPRSS2 protease encoding pCAGGS-TMPRSS2 plasmid were provided by S. Pöhlmann and M. Hoffman from the German Primate Center (Leibniz Institute for Primate Research). The lentiviral packaging plasmid p8.91 and firefly luciferase reporter plasmid pCSFLW were used for pseudotype production as previously described.^22^

Human embryonic kidney (HEK293T/17), human epithelial colorectal adenocarcinoma (Caco-2; ATCC:HTB-37), human fetal lung fibroblast-like (MRC-5; ATCC:HTB-37), monkey epithelial kidney cells (LLC-MK2; ATCC:CCL7), mink epithelial lung cells (Mv1Lu; ATCC:CCL64) and human adenocarcinoma alveolar basal epithelial (A549; ATCC: CRM-CCL-185) cells were maintained using Dulbecco’s Modified Eagle Medium (DMEM; GIBCO BRL, Paisley, UK) supplemented with 10% fetal bovine serum (FBS; Invitrogen, UK) and 1% penicillin/streptomycin (P/S; Sigma, Gillingham, UK). A549 clone 8 cells stably expressing ACE2 alone or ACE2 and TMPRSS2 were purchased from NIBSC (reference-codes: 101005 and 101006) and maintained as described^23^. All cell lines were grown in 37°C and in a humidified atmosphere of 5% CO_2_.

### Pseudotype virus production

Pseudotypes expressing the spike glycoproteins of NL63, 229E, SARS-CoV-2, VSV (vesicular stomatitis virus)-G or no (Δ-env) glycoprotein were produced as described.^24^ Briefly, the lentiviral packaging plasmid p8.91, the pCSFLW firefly luciferase vector, the pcDNA3.1^+^ expression plasmids for spike proteins were co-transfected using Fugene (Promega, Madison, WI, USA) into HEK293T/17 cells. Filtered supernatants were collected 48-72 h post-transfection. Twofold serial dilutions of PV-containing supernatant were performed as previously described using 96-well plates.^22^ Plates were incubated for 24-48 h, after which 50 µl Bright-Glo substrate (Promega, Southampton, UK) was added. Luciferase readings were conducted with a luminometer (GLOMAX™, Promega, Southampton, UK) after a 5-min incubation. Data were normalised using Δ-env and cell-only measurements and expressed as relative luminescence units (RLU) per ml.

### Virus infections, enzalutamide treatment and virus titration

All work involving the use of viruses was performed within BSL-2 laboratory under Health and Safety guidelines of the University of Essex. The HCoV-NL63 reference strain (isolate Amsterdam 1 [Ams-001]) was a kind gift from Dr. Lia van der Hoek and the OC43 and 229E strains were obtained from ATCC (VR-1558™ and VR-740™ respectively). The viruses were propagated in LLC-MK2, Mv1Lu and MRC-5 cells respectively. For antiandrogen treatment, cells were pre-treated with enzalutamide (ENZA, Stratech Scientific Ltd, UK; 1 μg/ml) 48-72h prior to infection. For infection assays, MRC-5 or Caco-2 cells were seeded in 12-well plates and incubated at 37°C with 5% CO_2_, until they reached 80% confluency. Unless otherwise specified, cells were infected for 2 hours with 229E, OC43 and NL63 respectively at a multiplicity of infection (MOI: 1). Media were then removed and replaced with DMEM and 2% heat-inactivated FBS, 1% P/S at 33°C for 2 days. The viruses released into the supernatant were then harvested and quantified by calculation of a 50% tissue culture infective dose per ml (TCID_50_/ml) according to the Reed–Muench formula and qPCR. Camostat mesylate (TMPRSS2 inhibitor) and E-64d (endosomal protease inhibitor) were obtained from MilliporeSigma (Bedford, UK).

### Real-Time Quantitative Reverse Transcription PCR (qRT-PCR)

All qRT-PCR analyses were conducted using the CFX96 Real-time PCR system (Biorad, Hemel Hempstead, UK). Viral RNA from the cell supernatant was extracted with TRIzol™ Reagent (Fisher Scientific, Leicestershire, UK) as per the manufacturer instructions. For OC43, NL63 and 229E nucleocapsid (*N*) gene PCR detection, a standard curve was generated using RNA dilutions of known copy number (ATCC standards: VR-1558DQ, VR-3263SD, VR-740DQ respectively) to allow absolute quantification of *N* copies from Ct values as previously described.^25^ Cell RNA isolation and RT-qPCR were performed using the PowerUp™ SYBR™ Green Master Mix (Fisher Scientific, Leicestershire, UK), following previously published procedures.^26,27^ Primer sequences for GAPDH, ACE2, TMPRSS2, Cathepsin-B, and Cathepsin-L were obtained from the literature.^9,28^ The quantitative gene expression data were normalised to the housekeeping gene GAPDH and compared with mock controls using plasmids containing the target genes as standards. Expression levels were quantified as gene copies per 1 μg RNA. All samples were tested in triplicate to ensure reproducibility.

### RNA-sequencing of nasal airway epithelial cells infected with HCoV

Air–liquid interface human nasal airway epithelial cells (HAE, MucilAir™; pool of 14 donors, catalog no. EP02MP; age and gender of donors are listed in supplemental material 1) were purchased from Epithelix and maintained in Mucilair cell culture medium (Epithelix, Geneva, Switzerland). HAE were kept at 5% CO_2_, 37°C. Briefly, before infection HAE were washed with serum-free media to remove mucus and debris. Cells were infected with 200 μL of virus-containing (HCoV-229E and -OC43) serum-free DMEM (MOI: 0.01) and incubated at 33°C for 1 h. Inoculum was then removed, and cells were washed twice. Time points were taken by adding 200 μL of serum-free DMEM and incubating for 10 mins and 37°C before removal and titration.

The cells were harvested at 24h and 72h for RNA using the RNeasy Mini Plus kit (Qiagen). Approximately 1 µg of RNA was used for the construction of sequencing libraries. The mRNA library preparation (poly-A enrichment) and sequencing were performed by Novogene following the manufacturer’s protocol. The sequencing data (FASTQ raw files) were imported into Partek Flow (version 10.0, build 10.0.23.0531; Partek Inc., St. Louis, MO, USA) for quality control and further processing. Paired-end reads were trimmed based on the Phred quality score threshold of >20 and aligned to the human genome (hg38) using the STAR-2.7.8a aligner. Quantification of gene expression was performed using the transcript model Ensembl Transcripts release 104 v2, and the data were subsequently normalised by the counts per million (CPM) method. Differential expressed genes were identified using the DESeq2 package. Genes were considered differentially expressed if they met the following criteria: p-value < 0.05, false discovery rate (FDR) < 0.05, and a fold change ≥ 2 or ≤ -2.

### Luciferase assay

MRC-5 cells were seeded at 1×10^4^ cells/well in hormone-depleted DMEM media in 96-well plates. After 24 hours, cells were co-transfected with TAT-GRE-EIB-LUC (500 ng/μl), pSVAR (100 ng/μl), and renilla (100 ng/μl) using FuGENE HD transfection reagent. At 48 hours, cells were either left untreated, or treated with 1 nM Mibolerone, and/or infected with OC43 or 229E (MOI 1) for 2 hours at 72 hours and lysed at 98 hours using 25 μl Dual-Glo® Luciferase Assay System (Promega). Luminescence was measured, and renilla luminescence was used for normalization with FLUOstar® Omega plate reader (BMG Labtech, Germany).

### Phylogenetic analysis

The amino acid sequences of coronavirus spike orthologues were subjected to multiple alignment using CLC Workbench 7 (CLC Bio, Qiagen, Aarhus, Denmark). Protein sequence NCBI reference sequences for the indicated viruses are as follows: SARS-CoV-2 (YP_009724390.1), 229E (QNT54842.1), NL63 (ANU06084.1), OC43 (CAA83660.1) and HKU1 (UJH59888.1).

### Data analyses

All analyses and graphs were performed using GraphPad Prism software version 9.2.0 (GraphPad, La Jolla, CA, US) and R, version 4.0.3 (R Foundation for Statistical computing). UpSet plots were generated using the R package UpSetR v1.4.0 and heat maps were generated using the Broad Institute’s Morpheus software (https://software.broadinstitute.org/morpheus/). The single-cell RNA sequencing dataset PRJNA789548 was re-processed, examined, and visualised using Cellenics® community instance (https://scp.biomage.net/) hosted by Biomage (https://biomage.net/). All data are presented as means with standard deviation (SD). Statistical analysis was carried out where possible to determine whether data was significant. Shapiro-Wilk normality tests were performed and based on the results we performed one/two-way ANOVA tests with Tukey’s multiple comparison tests. p<0.05 was considered significant unless otherwise stated.

### Data access

New high-throughput data generated in this study have been submitted to the NCBI Gene Expression Omnibus (GEO) (http://www.ncbi.nlm.nih.gov/geo/) under accession number GSE238079.

### Results and discussions

Research efforts are ongoing to develop vaccines and treatments for seasonal coronaviruses. Certain drugs that have been originally developed for other viral infections, such as interferons, ribavirin, and mycophenolic acid have shown promising outcomes against HCoV in clinical trials.^29^ Enzalutamide is a nonsteroidal antiandrogen medication that competitively hinders the interaction between androgens and the androgen receptor (AR), a transcription factor regulated by ligands.^30^ This action prevents the activation of genes that are under the control of AR, which is not only expressed in the prostate but also in the lungs.^31^ Additionally, enzalutamide has been shown to down-regulate TMPRSS2 expression, a type II transmembrane serine protease that cleaves the spike protein of SARS-CoV-2, preventing the virus from entering lung cells.^9^ This mechanism of action suggests that enzalutamide may have potential as a treatment option for COVID-19 and other viral infections that rely on TMPRSS2 for cell entry.

### The TMPRSS2 cleavage site of the spike protein is highly conserved among all HCoV

Viral attachment, fusion and cell entry of all CoV are mediated by their surface envelope spike (S) glycoprotein, a highly glycosylated type I fusion protein.^17,19,21^ The organisation of the ectodomain of the S protein in two subunits is shared in CoV: an N-terminal subunit named S1 that is responsible for binding to a cell surface receptor and a C-terminal S2 domain responsible for fusion of the viral and the cell membranes.^17^ The binding between S1 and a host cell receptor leads to conformational changes in the S protein, resulting in the fusion between virus and cell membranes and the release of viral nucleocapsid into the host cell cytoplasm.^17^ Many host cell proteases have been found to mediate the activation of S, including furin, cathepsin L, and trypsin-like serine proteases such as TMPRSS2, TMPRSS4, TMPRSS11A, and TMPRSS11D.^32,33^ Among them, TMPRSS2 and furin play a key role in the proteolytic activation of CoV and other viruses.^32^ Furin is expressed in the Golgi apparatus of most cells and cleaves polybasic or multi-basic sites of viral proteins.^32^ SARS-CoV-2, HKU1 and OC43 carry a functional furin cleavage site at the S1/S2 junction site (PRRAR↓ or NRRSRR↓; Figure 1A). TMPRSS2 is expressed in the human nasal, oral, and ocular mucosal surfaces and cleaves at a single arginine or lysine residue (R/K↓) at the S2’ site and plays a key role in triggering viral fusion for SARS-CoV-2 and other CoV.^33^ Amino acid sequence alignments of S orthologues illustrates that the TMPRSS2 cleavage site is well conserved among all four HCoV (Figure 1a).

**Figure 1:**
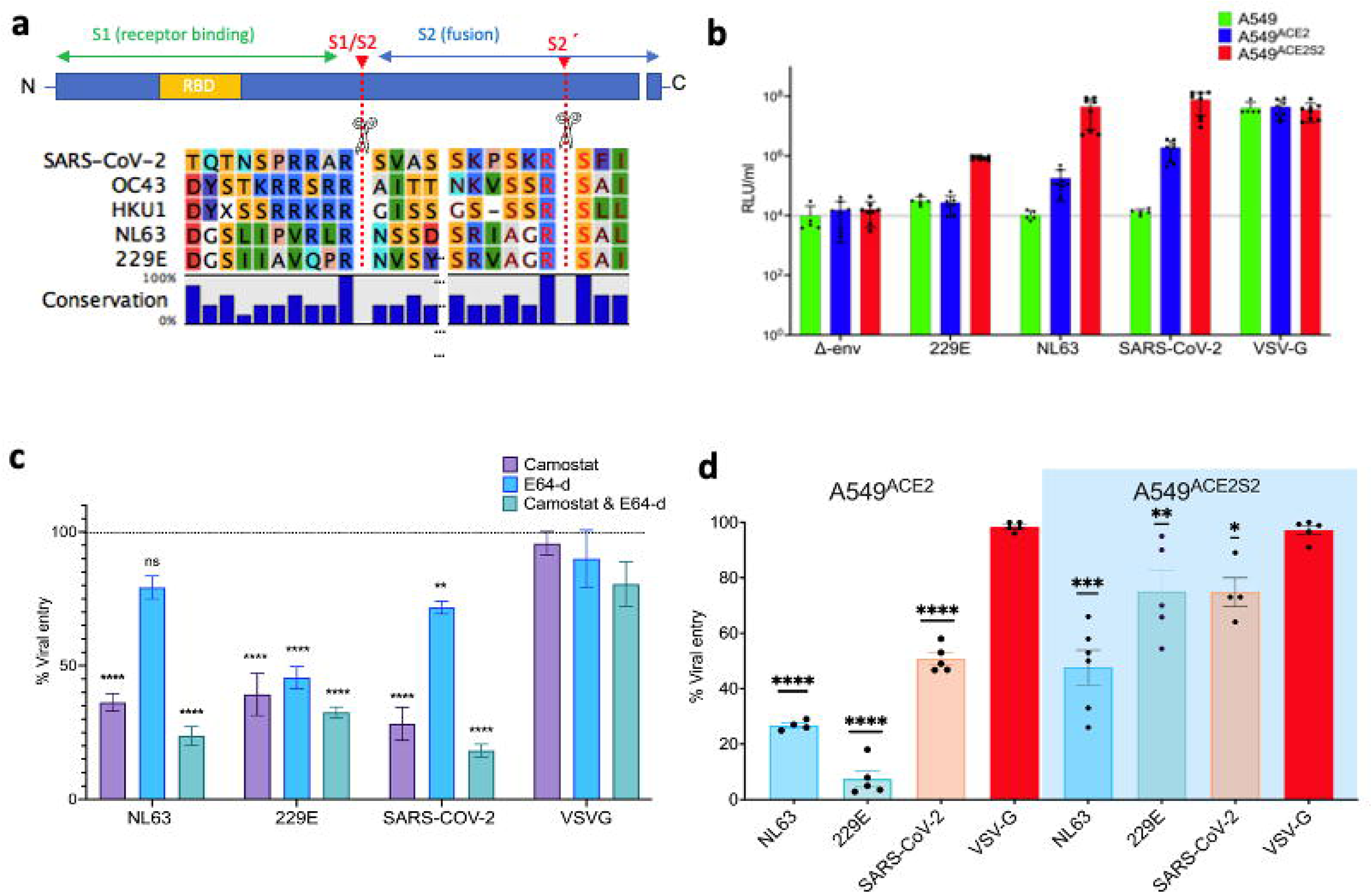
(**a**) Amino acid sequence alignment of S proteins from various human coronaviruses (HCoV) and SARS-CoV-2 at the S1/S2 and S2’ cleavage site. The receptor binding domain (RBD), spike glycoprotein subunits (S1 and S2), and S1/S2 and S2’ cleavage sites are labeled. **(b)** Transduction efficiencies (mean ± SEM, *n*=8) of Δ-env, NL63, 229E, SARS-CoV-2 S, and VSV-G pseudoviruses in A549, A549^ACE2^, and A549^ACE2S2^ cells at 48 h post transduction, as measured by luciferase activity and expressed as relative luminescence units (RLU/ml). (**c**) Percent pseudovirus (PV) transduction levels (mean ± SEM, *n*=3) in A549^ACE2S2^ cells pre-treated with camostat (50 μM) and/or E64-D (25 μM) inhibitors. Statistical significance was determined by two-way ANOVA and Tukey’s post hoc tests (\*\**p*=0.005, \*\*\*\**p*<0.0005, ns: not significant). (**d**) Percent PV transduction levels (mean ± SEM, *n*=6) in A549^ACE2^ and A549^ACE2S2^ cells pre-treated with enzalutamide (1 μg/ml) for 72 h. Statistical significance was determined by one-way ANOVA and Tukey’s post hoc tests (\**p*=0.05, \*\**p*=0.005, \*\*\**p*=0.0005, ****p<0.0001).

### Overexpression of TMPRSS2 facilitates cell entry of NL63 and 229E pseudotypes in A549 cells expressing ACE2 and TMPRSS2

To examine the functional significance of TMPRSS2 in the cell entry of HCoV, we employed the A549 cell line, derived from human lung adenocarcinoma, which is known to be androgen responsive.^31^ A549 cells express low constitutive levels of ACE2 and TMPRSS2 and thus are poorly permissive to infection by NL63, SARS-CoV-2 or spike pseudotyped lentiviral particles of HCoV and SARS-CoV-2 [in-house data] ^23^. For our study, we used two sublines of A549 cells expressing either ACE2 alone (A549^ACE2^) or both ACE2 and TMPRSS2 (A549^ACE2S2^).^23^ In our investigation into NL63 and 229E virus cell entry, we generated lentivirus pseudotypes (PV) incorporating their respective spike glycoproteins; however, we encountered challenges in producing PV for OC43, as the levels of OC43 lentiviruses were low (Supplemental material 2). PV carrying heterologous envelope proteins have played a pivotal role in the discovery of viral receptors^34^ and have proven valuable in evaluating the tropism of HCoV^24^. PV bearing SARS-CoV-2 S-and Vesicular stomatitis virus G-glycoproteins (VSV-G) served as positive controls; and PV without envelope protein (Δ-env) as negative controls. A549, A549^ACE2^ and A549^ACE2S2^ were transduced with the PV for 48h and viral entry was assayed *via* the expression of a PV-encoded luciferase reporter. Our data (Figure 1b) show that A549 were not susceptible to NL63- and SARS-CoV-2-PV but susceptible to VSV-G-PV and marginally susceptible to 229E-PV. A549^ACE2^ were more susceptible to NL63- and SARS-CoV-2-PV (by 30 and 140-fold transduction increase respectively) compared to wild type A549 as expected since ACE2 is the receptor for both NL63 and SARS-CoV-2. Conversely, A549^ACE2S2^ were more susceptible to all three PV compared to A549 (> 80-fold for each of them) suggesting that exogenous TMPRSS2 enhances cell entry of the viruses in line with published work.^35^

### Inhibition of TMPRSS2 by camostat mesylate reduces cell entry of NL63- and 229E-PV

To further investigate the significance of TMPRSS2 in the cell entry of HCoV, we treated A549^ACE2S2^ cells before PV transduction with the drugs camostat and E64-d, which are known to inhibit the activity of TMPRSS2 and cathepsin-L, respectively. Our results (Figure 1c) indicate that camostat reduced the entry of NL63, 229E, and SARS-CoV-2 PV by 74% (*p*<0.0001), 61% (*p*<0.0001), and 72% (*p*<0.0001), respectively. On the other hand, E64-d reduced the entry of NL63 PV by 21% (*p*=0.3067), 229E by 55% (*p*<0.0001), and SARS-CoV-2 by 29% (*p*=0.0026). When both E64-d and camostat were combined, we observed a further reduction in the entry of NL63, 229E, and SARS-CoV-2 by 87% (*p*<0.0001), 68% (*p*<0.0001), and 82% (*p*<0.0001), respectively. The positive control VSV-G did not show any significant reduction. The results suggest that A549^ACE2S2^ cells have an operational TMPRSS2-dependent pathway for the entry of NL63, 229E, and SARS-CoV-2 PV. The combined use of camostat and E64-d resulted in a more substantial reduction in viral entry, indicating the possible roles of both TMPRSS2 and cathepsin-L in facilitating the entry of these viruses. The smaller reduction in entry observed after treatment with E64-d alone suggests that cathepsin-L may play a less significant role in the entry of these viruses compared to TMPRSS2 in these cells. It’s possible that when TMPRSS2 levels increase in cells that usually have low levels of this protease, the infection may no longer depend on the cathepsin L pathway.^36^ Taken together, these findings suggest that HCoV, similar to SARS-CoV-2, can enter host cells through discrete, non-overlapping mechanisms and underscore the critical role of TMPRSS2 in the selection of entry pathway in line with previous research.^18–21^.

### Enzalutamide reduces cell entry of 229E and NL63-PV in A549^ACE2^ and A549^ACE2S2^ cells

To investigate the role of enzalutamide (ENZA) in inhibiting cell entry of NL63, 229E, and SARS-CoV- 2, we used PVs to transduce A549^ACE2^ and A549^ACE2S2^ cells, which were pretreated with ENZA for 72 hours. In A549^ACE2^ cells, ENZA significantly reduced viral entry of PV NL63, 229E, and SARS-CoV-2 by approximately 74%, 85%, and 50%, respectively (Figure 1d). To evaluate the potential influence of hormone levels on the observed outcomes, the experiments were replicated using hormone-depleted (charcoal stripped) media, yielding consistent results (Supplemental material 3). To further explore the contribution of TMPRSS2 in viral entry, we conducted an experiment using A549^ACE2S2^ cells. In these cells, the expression of exogenous ACE2 and TMPRSS2 were insensitive to androgen signaling, as the genes are under the control of the non-androgen responsive EF1a promoter.^23^ The results showed that ENZA partially reduced cell entry by approximately 60, 55, and 31% for NL63, 229E, and SARS-CoV-2, respectively. This suggests that when TMPRSS2 expression is regulated independently of androgen’s receptors, the effect of ENZA is lessened. Furthermore, ENZA did not impact the entry of VSV-G PV, indicating that ENZA’s effects are specific to HCoV and SARS-CoV-2 entry mechanisms. Additionally, we demonstrated that ENZA does not affect cell entry driven by VSV-G PV, indicating that the effects of the drug are specific for HCoV and SARS-CoV-2 mechanism of cell entry. Therefore, the partial reduction in viral entry in A549 cells stably expressing ACE2 alone and/or with TMPRSS2 by ENZA cannot be attributed solely to its influence on androgen signaling *via* TMPRSS2 regulation. This suggests that ENZA might have additional mechanisms of action that interfere with viral entry. Other androgen-responsive factors such as neuropilin-1, furin, TMPRSS4, and cathepsin G might also be involved in viral entry.^37–39^

### Enzalutamide reduces replication of authentic 229E and NL63 in TMPRSS2- expressing and non-expressing cells but not OC43

To confirm the results obtained with pseudotyped viruses, we quantified viral particle levels in the supernatant of Caco-2 and MRC-5 cells following 72 hours of ENZA treatment and subsequent infection with authentic 229E, OC43, and NL63 viruses, respectively (Figure 2a).We observed a significant reduction in viral copies from 2.79 x 10^7^ to 7.4 x 10^6^ for NL63 and from 3.54 x 10^6^ to 5.8 x 10^5^ for 229E, further confirming the inhibitory effect of ENZA on viral entry. Interestingly, enzalutamide did not have an effect against OC43.

**Figure 2:**
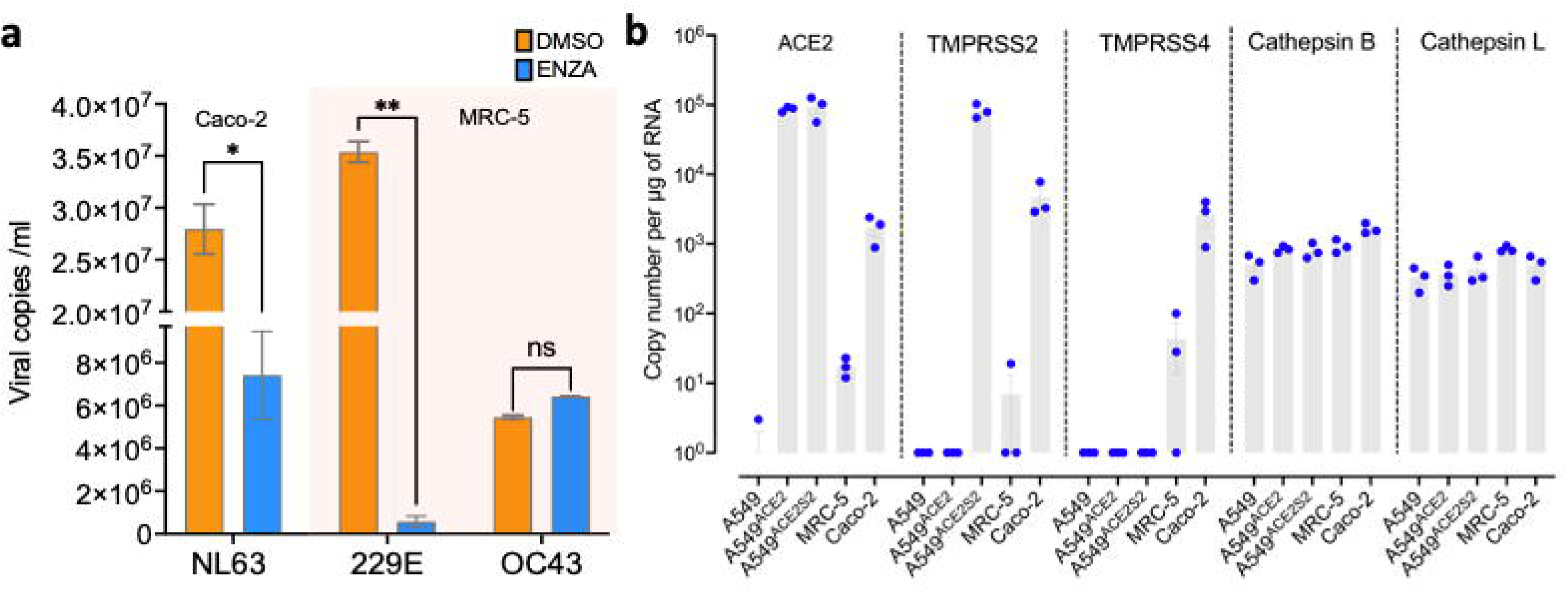
(**a**) Quantification of viral RNA copies in cell culture supernatant after pre-treatment of MRC-5 and Caco-2 cells with enzalutamide (1 μg/ml) for 72 h and infection with 229E and NL63 viruses (MOI: 1), respectively, as determined by qRT-PCR for the nucleocapsid (*N*) gene. DMSO-treated cells were used as controls in each cell line. Analysis was performed in two independent experiments. Statistical significance was determined by two-way ANOVA and Šidák’s post hoc tests (\**p*=0.05). (**b**) Quantification of ACE2 and proteases (TMPRSS2, TMPRSS4, Cathepsin B and L) copy numbers per μg of total cellular RNA in the examined cell lines was carried out using qRT-PCR analysis, with plasmids serving as reference standards for precise calculation.

To investigate the mechanism of action of enzalutamide, we assessed the mRNA expression levels of ACE2 and proteases involved in the cellular entry of coronaviruses by RT-qPCR (Figure 2b). Our findings demonstrate that NL63-permissive Caco-2 cells express TMPRSS2, TMPRSS4, and ACE2, while 229E-and OC43-permissive MRC-5 cells exhibit no constitutive expression of TMPRSS2, TMPRSS4, and ACE2. In contrast, A549 cells required lentiviral transduction to establish stable ACE2 and TMPRSS2 expression, to render them susceptible to 229E/NL63 and SARS-CoV-2 pseudoviruses. Conversely, non-androgen responsive cathepsins B and L exhibited constitutive expression across all tested cell lines. Taken together these results suggest that enzalutamide may also exert its effect via non-TMPRSS2-mediated mechanisms that inhibit viral entry and possibly replication. Enzalutamide’s antiviral effects may be attributed to its impact on other proteases or cellular factors that participate in the entry or replication of coronaviruses, such as cathepsins or furin, or could be a result of off-target effects, and not related to its primary function as an androgen receptor inhibitor. Further research is necessary to elucidate the exact mechanism by which enzalutamide is exerting its antiviral effects.

### Distinct androgen signaling responses to HCoV infections in primary human nasal airway cells

Previous studies have demonstrated that viruses can act as noncellular positive coregulators for androgen receptor (AR).^38,40,41^. The AR governs the expression of target genes by binding to specific cis-regulatory elements, a process primarily facilitated by its DNA-binding domain (DBD). Upon binding with androgen, the AR translocates into the nucleus, where it forms homodimers and directly interacts with DNA, often at the consensus AR-binding motif known as canonical AREs.^31^

In the context of our investigation, we aimed to assess the influence of HCoV on AR activity, providing insights into the varied effects of enzalutamide across different HCoV. To achieve this, we conducted RNA-sequencing analysis (RNA-seq) on air-liquid interface nasal epithelial cells (HAE) infected with two distinct strains: HCoV 229E, belonging to the alpha lineage and HCoV OC43, representing the beta lineage. Notably, our choice to study both strains was deliberate; enzalutamide demonstrated efficacy against 229E but not OC43, prompting a focused exploration of these specific HCoV. The HAE cells were derived from a pool of 14 human donors, ensuring representation of both genders in the study. These primary cells faithfully replicate normal airway biology, featuring pseudostratified mucociliated differentiation and organotypic cell types.^42^

Our primary objective was to uncover enrichment patterns between gene sets influenced by the virus and genes responsive to androgen. We assessed AR activity through three well-established gene signatures^43–45^, revealing significant disparities in how AR-regulated genes responded to the two viruses, as illustrated in Figure 3a-b. Specifically, 229E induced changes in a subset of AR-regulated transcripts (22 and 135 at 24hpi and 72hpi respectively), while OC43 affected only a limited number of such transcripts (8 and 21 at 24hpi and 72hpi respectively) (Fig. 3a-b). The full dataset of androgen-responsive gene expression is contained in Supplemental material 4. Our attempts to detect upregulation of the *AR* through RNA-seq reads and immunoblotting yielded inconclusive results, likely due to low detection sensitivity and cellular heterogeneity. Moreover, the mRNA expression of *AR* gene alone, quantified through RNA-seq, was not significantly regulated in virus-infected samples when compared to uninfected samples. This led us to hypothesize that AR might still be active, albeit with restricted expression in specific nasal airway cell types. To gain further insights, we reanalysed publicly available single-cell RNA-seq data of HAE, revealing active AR expression that is relatively enriched in epithelial and airway goblet cells compared to other nasal epithelial cell types (Fig. 3c). Together, these findings indicate a significant influence of 229E on global androgen signaling even in primary cells with low AR expression.

**Figure 3:**
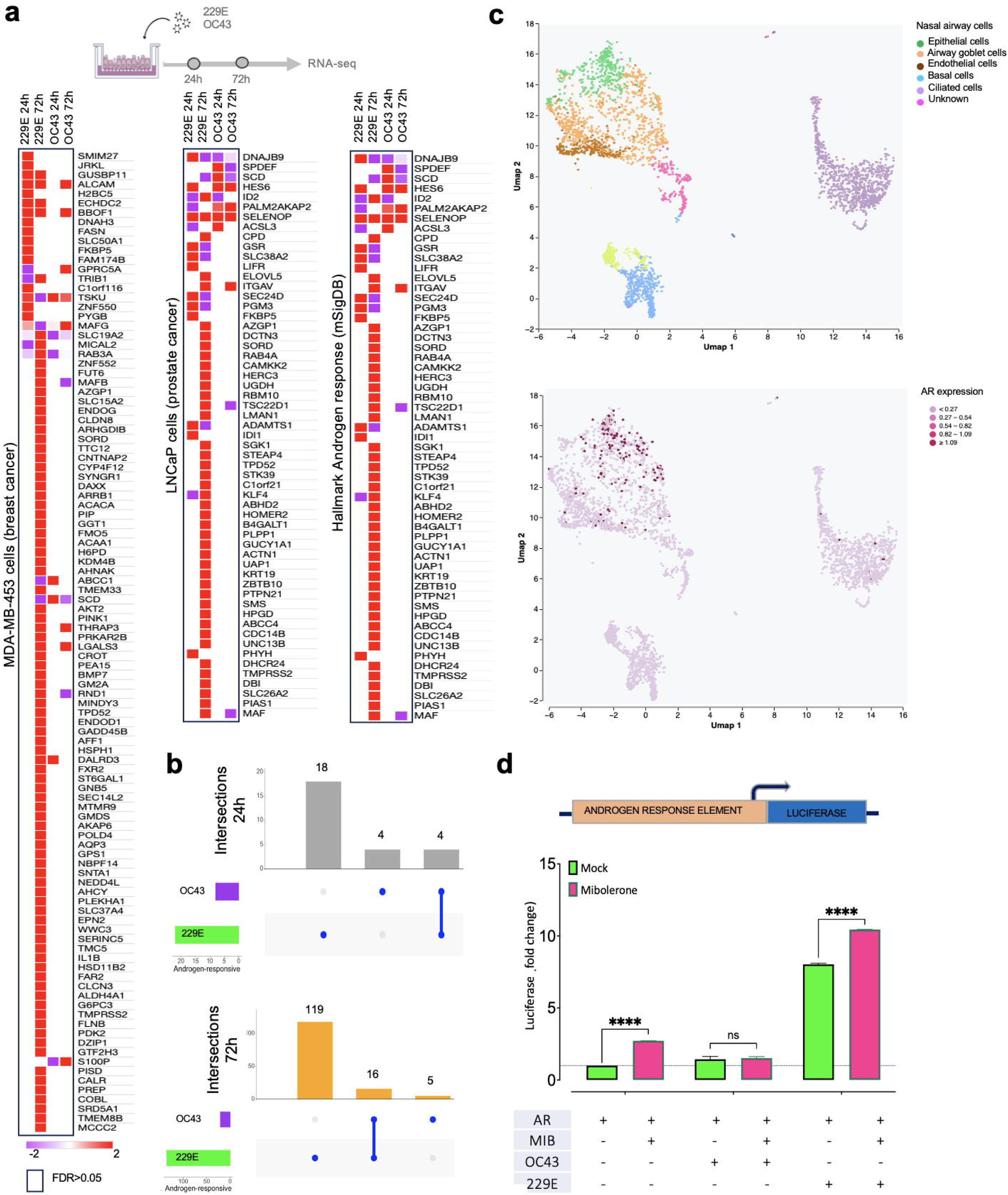
Distinct androgen signaling responses to viral infections: (**a**) Primary human nasal airway cells were infected with human coronaviruses OC43 and 229E, and the differential gene expression analysis at 24 and 72h post infection was performed using bulk RNA-sequencing. The heatmaps shows relative transcript expression of androgen-responsive transcripts within three datasets: MD-SB1 breast cancer cells^44^, LNCaP cells^43^ and the hallmark androgen response gene set M5908 (mSiGdb)^45^. (**b**) Upset plots illustrating the overlap of androgen-responsive transcripts modulated by 229E and OC43 viruses at 24 hours (upper panel) and 72 hours (lower panel) post-infection. Bar plots represent the count of androgen-responsive transcripts, while dots connected by lines depict the transcripts shared between the two viruses. (**c**) Single-cell RNA-seq analysis of AR expression: (Upper) UMAP visualization of the cell populations nasal HAE samples UMAP plot of the single cell RNA-seq clustering from the bronchoalveolar samples from COVID-19 and healthy control individuals. A total of 34 clusters were found in this sample of 6 severe and 3 mild COVID-19 patients as well as 3 healthy controls. Each cluster is colored differently and depicted by a number. Each cluster is colored differently and depicted by a number. (lower) UMAP visualization of the AR distribution in HAE. (**d**) Androgen response element (ARE) luciferase assay: MRC-5 cells were co-transfected with an AR expression vector and an ARE-luciferase reporter. Cells were infected with OC43 and 229E viruses, and luciferase activity was measured. The bar plot represents the luciferase activity relative to the control. Statistical significance was determined by one-way ANOVA and Tukey’s post hoc tests (\**p*=0.05, \*\**p*=0.005, \*\*\**p*=0.0005, ****p<0.0001).

To investigate whether the elevated expression of the androgen-induced core genes can be attributed to a direct effect of the virus on the androgen response element (ARE), we employed a luciferase construct controlled by the ARE (Fig. 3d). MRC-5 cells which exhibit low androgen receptor expression and were maintained in androgen-depleted media, were infected with OC43 and 229E viruses, and co-transfected with an AR expression vector and an ARE-luciferase reporter. MRC-5 cells infected with 229E exhibited strong luciferase activity both when exposed to mibolerone (a synthetic androgen) and when not exposed to it. This finding highlights a significant increase in luciferase activity during 229E infection, irrespective of mibolerone presence, strongly indicating a direct association between viral infection and androgen signaling. Interestingly, OC43 infection had an opposing effect, inhibiting the activation of ARE with/-out mibolerone, potentially explaining the lack of response to the anti-androgen enzalutamide in OC43-infected cells.

In conclusion, our study highlights the intricate relationship between viral infections and host hormone pathways, specifically androgen signaling, in cell lines with low androgen receptor expression. It suggests that viruses can manipulate androgen signaling, either by mimicking or antagonizing the AR pathway. These insights pave the way for advancements in managing viral infections and developing therapeutic interventions. In this context, enzalutamide’s ability to block the androgen receptor may impede the entry and replication of 229E and NL63 coronaviruses through both TMPRSS2-dependent and TMPRSS2-independent pathways. Given the absence of approved antiviral drugs for seasonal coronaviruses, our research presents a potential avenue for novel therapeutic options centered around antiandrogens. Further exploration, particularly through clinical trials, is essential to validate these findings and translate them into effective antiviral strategies.

### Funding

Funding was provided by the University of Essex COVID-19 Rapid and Agile and the Faculty of Science and Health Research Innovation and Support Funds. Tukur Abdullahi is supported by the Nigerian Petroleum Technology Development Fund and the University of Essex.

### Author Contributions

O.D.O. performed experiments, analysed data and wrote the first draft of the manuscript; T.A., R.S. and E.S.G. performed experiments and analysed data; C.L.B, N.T. and G.N.B. provided reagents and edited the manuscript; E.S.G. conceptualised the study, supervised activities and edited the manuscript. All authors have read and agreed to the published version of the manuscript.

## Abbreviations

COVID-19: Coronavirus Disease 2019
SARS-CoV: Severe acute respiratory syndrome coronavirus
NL63: Human coronavirus NL63
229E: Human coronavirus 229E
OC43: Human coronavirus OC43
TMPRSS2: Transmembrane protease serine 2
ACE2: angiotensin-converting Enzyme 2
ANOVA: analysis of variance
GAPDH: Glyceraldehyde 3-phosphate Dehydrogenase
ATCC: American Type Culture Collection
MOI: Multiplicity of Infection
BSL-2: Biosafety level 2.

## Acknowledgments

We thank Tiffany Teoh and Veronica Mavrovouna for their technical assistance.

## Supplemental material

**Supplemental material 1:** List of donors used to generate the study’s nasal epithelial airway cells (Data were provided by Epithelix, Geneva, Switzerland).

**Supplemental material 2:** Figure illustrates the susceptibility of cells to OC43 pseudoviruses (PVs) that either express or lack the spike (S) glycoprotein, with varying levels of membrane (M) protein expression plasmid. Cell susceptibility is assessed through transduction titers (*n* = 8) in different cell lines. The luminescence-based measurement of cell entry is presented on a log10 scale as Relative Luminescence Units per milliliter (RLU/ml) and is visualised using box plots. PVs are compared to a negative control (Δ-env, grey bars), which helps establish the baseline of background luminescence signal in the target cells (103 – 104 RLU/ml). The data are presented as average means, with error bars indicating standard deviation (SD).

**Supplemental material 3:** Percent PV transduction levels (mean ± SEM, *n*=6) in A549^ACE2^ cells grown in charcoal-stripped serum and pre-treated with enzalutamide (1 μg/ml) for 72 h. Statistical significance was determined by one-way ANOVA and Tukey’s post hoc tests (****p<0.0001).

**Supplemental material 4:** Excel workbook displaying fold changes in androgen-responsive gene expression induced by NL63 and OC43 viruses in HAE cells. Expression profiles were cross-referenced with AR transcriptional regulation databases: MDA-MB453 breast cancer cells (Doane et al.)^42^, LNCaP prostate cancer cells (Nelson et al.)^43^, and Hallmark Androgen response (M5908, mSigDB)^44^, as shown in Figure 3a-b. Data depict virus-infected versus mock-infected cell comparisons in different worksheets representing distinct databases. Empty squares denote missing data due to low expression or limited statistical significance. Colors indicate upregulation (blue) or downregulation (red).

