## Supplementsl material 1-3 for "The antiviral potential of the antiandrogen enzalutamide and the viral-androgen interplay in seasonal coronaviruses": Supplemental material.pdf

**Supplemental material 1:** List of donors used to generate the study's nasal epithelial airway cells (Data were provided by Epithelix, Geneva, Switzerland).

| Batch number | Age | Gender |
|---|---|---|
| AB0870 | 77 | F |
| AB0866 | 54 | M |
| AB0863 | 57 | F |
| AB0825 | 65 | F |
| AB0824 | 69 | M |
| AB080901 | 24 | M |
| AB080701 | 61 | M |
| AB079401 | 53 | M |
| AB077401 | 38 | M |
| AB077101 | 23 | M |
| AB076700 | 71 | M |
| AB076001 | 45 | N/A |
| AB074501 | 45 | M |
| AB050701 | 29 | M |

**Supplemental material 2:** Figure illustrates the susceptibility of cells to OC43 pseudoviruses (PVs) that either express or lack the spike (S) glycoprotein, with varying levels of membrane (M) protein expression plasmid. Cell susceptibility is assessed through transduction titers (n = 8) in different cell lines. The luminescence-based measurement of cell entry is presented on a log10 scale as Relative Luminescence Units per milliliter (RLU/ml) and is visualized using box plots. PVs are compared to a negative control (Δ-env, grey bars), which helps establish the baseline of background luminescence signal in the target cells (103 – 104 RLU/ml). The data are presented as average means, with error bars indicating standard deviation (SD).

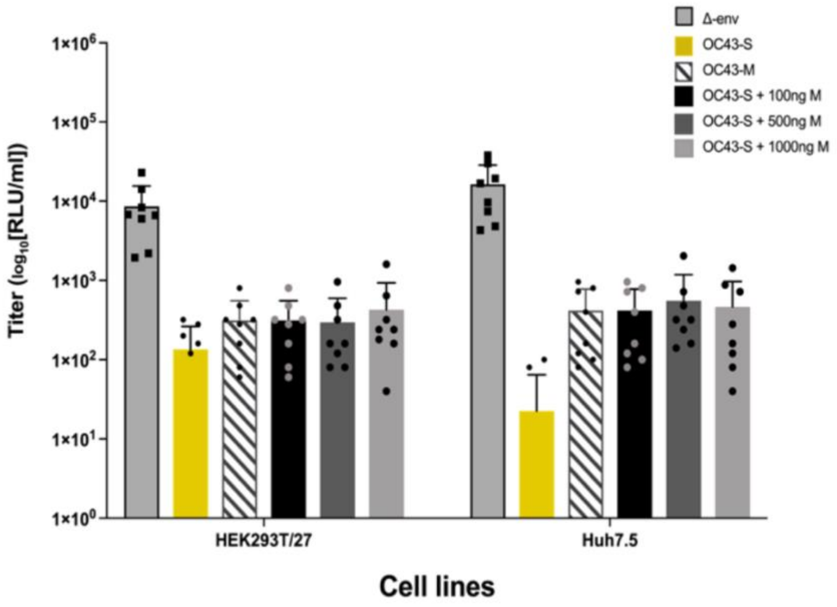

**Supplemental material 3:** Percent PV transduction levels (mean ± SEM, *n*=6) in A549[ACE2] cells grown in charcoal-stripped serum and pre-treated with enzalutamide (1 µg/ml) for 72 h. Statistical significance was determined by one-way ANOVA and Tukey's post hoc tests (****p<0.0001).

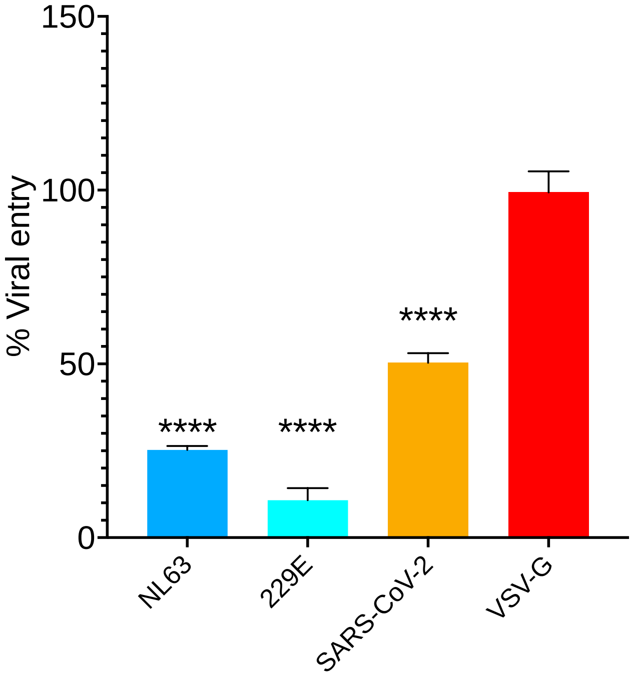
